# *TP53* minigene analysis of 161 sequence changes provides evidence for role of spatial constraint and regulatory elements on variant-induced splicing impact

**DOI:** 10.1101/2024.10.07.617118

**Authors:** Daffodil M. Canson, Inés Llinares-Burguet, Cristina Fortuno, Lara Sanoguera-Miralles, Elena Bueno-Martínez, Miguel de la Hoya, Amanda B. Spurdle, Eladio A. Velasco-Sampedro

## Abstract

Germline *TP53* genetic variants that disrupt splicing are implicated in hereditary cancer predisposition, while somatic variants contribute to tumorigenesis. We investigated the role of *TP53* splicing regulatory elements (SREs), including G-runs that act as intronic splicing enhancers, using exons 3 and 6 and their downstream introns as models. Minigene microdeletion assays revealed four SRE-rich intervals: c.573_598, c.618_641, c.653_669 and c.672+14_672+36. A diagnostically reported deletion c.655_670del, overlapping an SRE-rich interval, induced an in-frame transcript Δ(E6q21) from new donor site usage. Within intron 6, deletion of at least four G-runs led to 100% aberrant transcript expression. Additionally, assay results suggested a donor-to-branchpoint distance cutoff of <50 nt for complete splicing aberration due to spatial constraint, and >75 nt for low risk of splicing abnormality. Overall, splicing data for 134 single nucleotide variants (SNVs) and 27 deletions in *TP53* demonstrated that SRE-disrupting SNVs have weak splicing impact (up to 26% exon skipping), while deletions spanning multiple SREs can have profound splicing effects. Results also provide more data to inform splicing impact prediction for intronic deletions that shorten intron size.

## Introduction

Pre-mRNA splicing, the removal of introns followed by exon ligation to produce the mature mRNA, is a key step in the expression of most human genes. Alternative splicing plays a vital role in controlling gene expression and enhances the complexity of the transcriptome and proteome by allowing a single gene to produce multiple mRNA transcripts^1^. Constitutive and alternative pre-mRNA splicing are controlled by a wide array of factors and sequence motifs including the 5’ (donor) and the 3’ (acceptor) splice sites, the polypyrimidine tract (PPT), the branch point (BP), and splicing regulatory elements (SREs) such as exonic or intronic splicing enhancers (ESE/ISE) or exonic or intronic splicing silencers (ESS/ISS)^2^. *Trans*-acting RNA binding proteins (RBPs) known as splicing factors regulate exon inclusion, exclusion, or alternative splice site usage by binding to SREs within the pre-mRNA exons or their flanking introns^3^. The commonly recognized RBPs that target SREs include, but are not limited to, serine/arginine-rich (SR) proteins and heterogeneous nuclear ribonucleoproteins (hnRNPs)^4^. Splicing factors and SREs regulate splicing in a position- and context-dependent manner^4^. For example, RBPs of the hnRNP F/H gene family bind to sequences of three or more consecutive guanines (G-runs) that act as enhancers when located downstream of a donor site^5^, and conversely as silencers when located in the exon^6^. Genetic variants that affect splicing motifs and disrupt the binding of splicing factors can cause defective splicing resulting in abnormal mRNA transcripts, or modify the relative levels of alternatively spliced isoforms. Splicing alterations are causally associated with cancer susceptibility, development, progression, and prognosis^7,8^.

The p53 tumor suppressor exists in several alternative isoforms, making it a good model for the investigation of *cis*-regulatory elements and RBPs. The *TP53* gene is composed of 11 exons and two cryptic exons (9β and 9γ) within intron 9^9^. *TP53* alternative splicing, alternative promoter usage, and alternative initiation of translation generates at least 16 different p53 protein isoforms^10^. Several studies have shown that p53 isoforms are abnormally expressed in a wide array of cancers^9^. For example, elevated expression of the Δ40p53 isoform is associated with the aggressive triple negative breast cancer^11^. *TP53* intronic variants can affect the production of alternative p53 isoforms. Minigene experiments measuring the effect of two G-run deletions and site-directed mutagenesis of these G-runs in intron 3 showed increased levels of mRNA with intron 2 retention that encodes the Δ40p53 isoform^12^. Somatic variants at the intron 4 donor site are associated with overexpression of transcripts encoding the Δ133p53 isoforms in breast tumors^13^. Somatic variants can also produce additional aberrant isoforms such as the intron 6 acceptor site variants that activate an intronic cryptic acceptor, generating the truncated p53Ψ isoform that promotes metastasis^14^. Moreover, *TP53* rare germline variants that cause splicing aberrations are associated with Li Fraumeni Syndrome^15,16^, a hereditary multicancer predisposition syndrome characterized by early-onset cancers primarily breast cancer, bone and soft tissue sarcomas, brain tumors, and adrenocortical carcinomas^17^.

Previous studies on variant-induced *TP53* aberrant or alternative splicing have mostly focused on the donor and acceptor splice consensus motifs^18–20^. In this study, we aimed to investigate the splicing impact of variants located in *TP53* SRE-rich regions and predict the RBPs potentially targeting these *cis*-regulatory elements. We prioritized *TP53* exons 3 and 6, as they are suitable candidates for being regulated by SREs for two different reasons. First, the *TP53* exon 3 size (22 nt) falls into the category of “microexons” (<30 or 51-nt depending on the authors), which would require specialized regulatory factors involved in their recognition and inclusion^21,22^. Second, the weakness of exon 6 native donor site, as well as the presence of a strong intronic cryptic donor site 63 nt downstream, suggest a specific regulation that would promote recognition of the native donor site through SREs. Further, *TP53* introns 3 and 6 contain several G-rich sequences that represent putative binding motifs for splicing factors, such as hnRNP A/B and F/H that play a role in intron definition^23^. Notably, intron 3 contains G-quadruplex structures that control intron 2 excision in H1299 cells^12^.

In previous reports, we have shown that minigene analysis is a suitable approach for studying the splicing effect of variants in breast cancer susceptibility genes^24–28^. Here, we designed a splicing reporter minigene with *TP53* exons 2 to 9, where we experimentally tested for the presence of SRE-rich intervals by analyzing 23 different microdeletions in exons 3 and 6 and their downstream introns. We also tested the impact of four deletions detected in patients (reported in the ClinVar database) and 134 single nucleotide variants (SNVs) on *TP53* splicing.

## Materials and Methods

### Bioinformatics and databases

Variant data and alternative transcripts were annotated based on the *TP53* MANE Select transcript (NM_000546.6), composed of 11 exons that encodes the canonical p53α protein. Splicing events were described with a short descriptor combining the following symbols: Δ (skipping of exonic sequences), ▼ (inclusion of intronic sequences), E (exon), p (acceptor site shift) and q (donor site shift)^29,30^. When necessary, the number of deleted or inserted nucleotides is indicated. For example, ▼(E6q5) indicates the use of an intronic cryptic donor 5 nt downstream of exon 6, generating a 5-nt insertion into the mature mRNA.

*TP53* sequences spanning exon 3, intron 3, exon 6, and part of intron 6 (from c.672+1 to c.672+85) were analyzed with the following online *in silico* prediction tools for SREs: i) HEXplorer for mapping out the putative enhancer and silencer regions^31^; (ii) SpliceAid, a database of experimentally derived target RNA sequences of RBPs^32^; and (iii) DeepCLIP, a deep learning approach, which integrates RNA binding functional data to predict the presence of splicing factor motifs^33^. In addition to SRE mapping, these tools were used to predict the impact of deletions and SNVs on putative SRE motifs and on RBP binding. For HEXplorer analysis of SNVs, the delta HZEI score of 11-nt sequences containing the SNV in the middle was calculated; delta HZEI ≤ -5 (exonic) or ≥5 (intronic) was considered as SRE-disrupting^34^. A similar approach of analyzing 11-nt sequences was used to generate DeepCLIP predictions for SNVs. Cutoffs for DeepCLIP analysis were set as follows: RBP binding score >0.8 for putative RBP binding site in the wild type and variant sequences, and a decrease in binding score by 0.3 or greater for binding site disruption. RBPs considered as splicing factors that regulate splice site usage by binding to exonic or intronic motifs, and also listed in SpliceAid were included. RBPs that bind to both wild type and variant sequences were excluded.

Splicing predictions were done using SpliceAI v1.3.1^35^ with maximum distance of ±4999 nt flanking the variant, unmasked, and GRCh38 genome assembly. SpliceAI cutoff scores were ≥0.2 for variants predicted to be spliceogenic (i.e. to impact native splicing patterns) and ≤0.1 for variants predicted to be non-spliceogenic^36^. The strength of the acceptor and donor consensus motifs was predicted by MaxEntScan (MES)^37^. PhyloP^38^ was used to measure nucleotide conservation of intronic SNVs. Minor allele frequency of SNVs was obtained from the gnomAD v2.1.1 non-cancer dataset^39^. Clinical variants and their pathogenicity assertions were obtained from the ClinVar database^40^.

### Minigene construct design and site-directed mutagenesis

A 3487-bp insert, including exons 2 to 9, was designed in-house (**Supplementary Figure S1**) and then generated by gene synthesis (GeneArt, Thermo Fisher Scientific, Waltham, MA, USA). This fragment was cloned into the pSAD splicing vector between the restriction sites *Sac*I and *Eco*RI to obtain the minigene mgTP53_2-9 (**Figure 1A**), which was confirmed by sequencing (Macrogen, Madrid, Spain).

**Figure 1.**
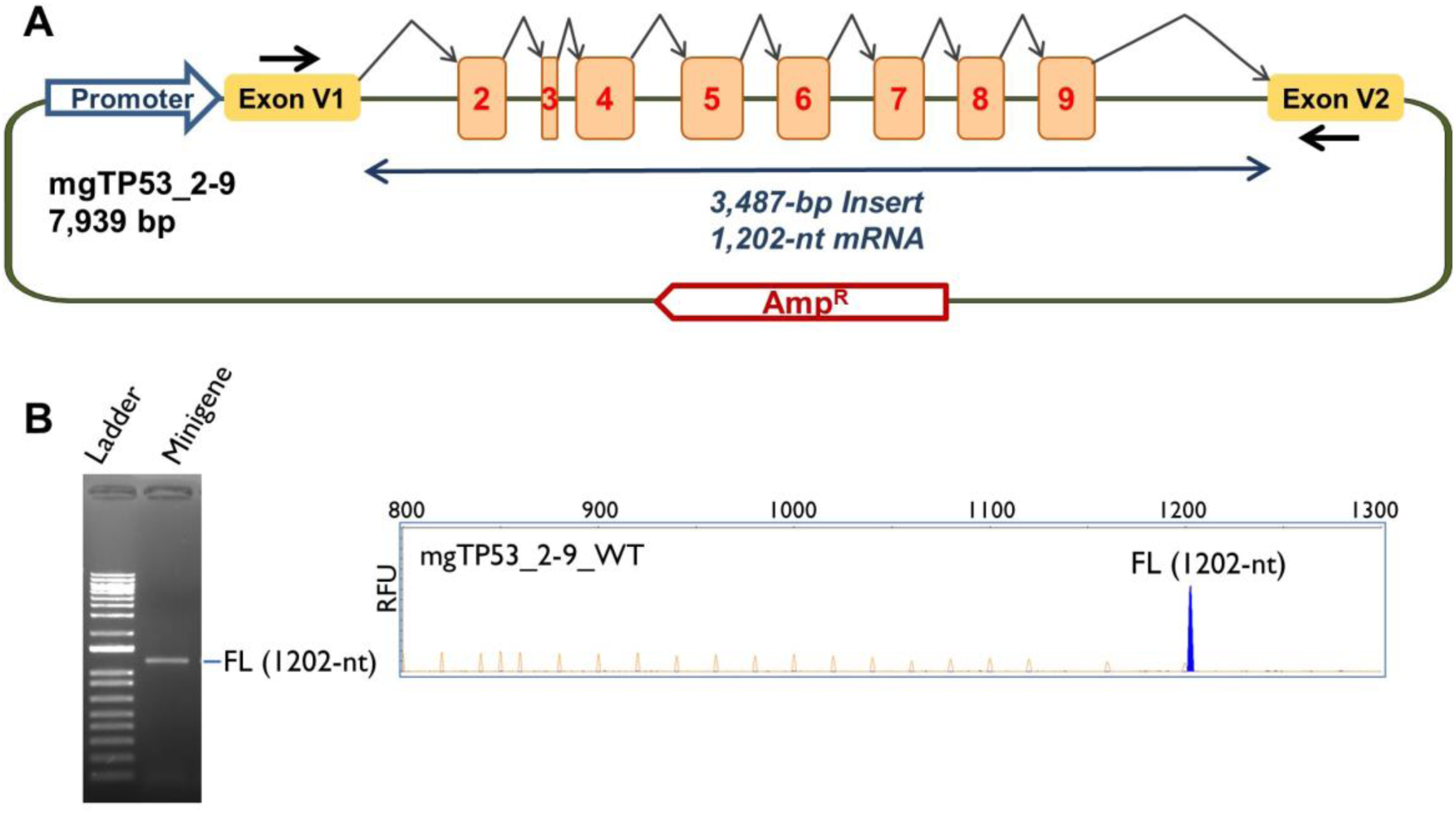
Structure and splicing assay of the *TP53* minigene with exons 2 to 9. **A)** Schematic representation of the minigene mgTP53_2-9 with exons 2 to 9 in numbered boxes. Black arrows indicate the location of vector-specific RT-PCR primers. **B)** Splicing assay of the WT minigene. RT-PCR products were analyzed by agarose gel electrophoresis (left) and fluorescent fragment electrophoresis (right). FL, expected minigene full-length transcript. The x-axis indicates size in bp and the y-axis represents Relative Fluorescence Units (RFU).

Microdeletions and candidate variants (**Supplementary Table S1**) were introduced into the wild type (WT) construct by site-directed mutagenesis using the QuikChange Lightning Kit (Agilent, Santa Clara, CA, USA). Mutant minigenes were confirmed by sequencing (Macrogen). Additionally, the G-rich *TP53* intron 3 was replaced with *ATM* intron 21 (as a control/reference intron which does not contain G-runs) (**Supplementary Figure S2**) by overlap extension PCR as previously described^41^.

### Splicing assays

#### Transfection and RNA isolation

Approximately 2 x10^5^ cells of the human breast cancer cell line SKBR3 (ATCC HTB-30) were grown in 0.5 mL of medium (Minimum Essential Medium -MEM-, 10% Fetal Bovine Serum, 1% nonessential amino acids, 2mM Glutamine and 1% Penicillin/Streptomycin; Sigma-Aldrich, St. Louis, MO, USA) in 4-well plates (Nunc, Roskilde, Denmark). SKBR3 cells were transiently transfected with 1 µg of the WT/mutant minigenes and 2µL of lipofectamine LTX (Life Technologies, Carlsbad, CA, USA).

In order to inhibit nonsense-mediated decay (NMD), a 4-hour incubation with 300 µg/ml of cycloheximide (Sigma-Aldrich, St. Louis, MO, USA) was carried out 48 hours after transfection. RNA Purification was performed using the Genematrix Universal RNA Purification Kit (EURx, Gdansk, Poland), with on-column DNAse I digestion.

To check splicing profile reproducibility, the WT and four variant constructs (c.76C>A, c.592G>T, c.655_670del, and [c.672+14_672+36del;c.672+39_672+46del]) were also tested in U2OS osteosarcoma and HeLa cell lines following the same protocols as above.

#### Reverse transcription polymerase chain reaction (RT-PCR)

A total of 400 ng of RNA were retrotranscribed with the RevertAid First Strand cDNA Synthesis Kit (Life Technologies), using the vector-specific primer RT-PSPL3-RV (5′-TGAGGAGTGAATTGGTCGAA-3′) and the manufacturer’s protocol.

The resulting cDNA was amplified with primers SD6-PSPL3_RT-FW (5’-TCACCTGGACAACCTCAAAG-3’) and RTpSAD-RV (CSIC Patent P201231427) using Platinum Taq DNA polymerase (Life Technologies) and the following cycling conditions: 94°C/2 min, 35 cycles x [94°C/30 s, 60°C/30 s, 72°C/(1 min/kb)], 1 cycle x [72°C/5 min]. RT-PCR products were sequenced by Macrogen. The expected size of the minigene full-length (FL) transcript was 1202-nt.

In order to quantify the relative proportions of each transcript, semi-quantitative fluorescent RT-PCRs (26 cycles) were carried out in triplicate using Platinum Taq DNA polymerase (Life Technologies) and the primers SD6-PSPL3_RT-FW and RTpSAD-RV (FAM-labelled). Fluorescent products were run with LIZ-1200 size standard at the Macrogen facility and analyzed using Peak Scanner_V1.0 (Life Technologies). Only peak heights ≥100 RFU (relative fluorescence units) were considered. Mean peak areas of each transcript and standard deviations were calculated.

## Results

The minigene construct mgTP53_2-9 (**Figure 1A**) produced the expected FL transcript of 1202-nt (V1 – *TP53* exons 2 to 9 – V2) without any alternative transcript when assayed in SKBR3 cells (**Figure 1B**), resembling the naturally occurring pattern found in breast and ovary (data not shown). Additionally, the splicing patterns produced from SKBR3, U2OS, and HeLa cells were identical on agarose gel, indicating reproducibility of splicing profile (**Supplementary Figure S3**). The SpliceVault^42^ 300K-RNA data and its sub-group of breast samples also showed a low frequency of alternative splicing events in *TP53* exons 2 to 9, supporting a very low level of naturally occurring alternative splicing in this region.

### Functional mapping of splicing regulatory elements

HEXplorer analysis revealed several putative SRE-rich regions in intron 3, intron 6, and exon 6 (**Supplementary Figure S4**). To determine if they play a role in splicing, we analyzed several intronic microdeletions spanning hnRNP clusters and putative KSRP, SF1, and TIA1 binding sites, and exon 6 microdeletions spanning putative ESE motifs (**Figure 2, Supplementary Table S2**). Microdeletions affecting putative SREs were designed such that none were predicted to create a new donor or acceptor motif. Concerning introns 3 and 6, we focused on six clusters (three in each intron) of hnRNP A1 and H motifs predicted by SpliceAid in the G-rich regions (**Figure 2A-B**). SpliceAid analysis of intron 3 also predicted the presence of binding sites for the splicing factors KSRP and SF1 that are involved in microexon recognition ^22^. Moreover, SpliceAid predicted a binding site in intron 6 for the TIA1 protein that is known to promote the usage of weak donor sites^43^. We excluded exon 3 from the microdeletion analysis because of its small size of 22 nt.

**Figure 2.**
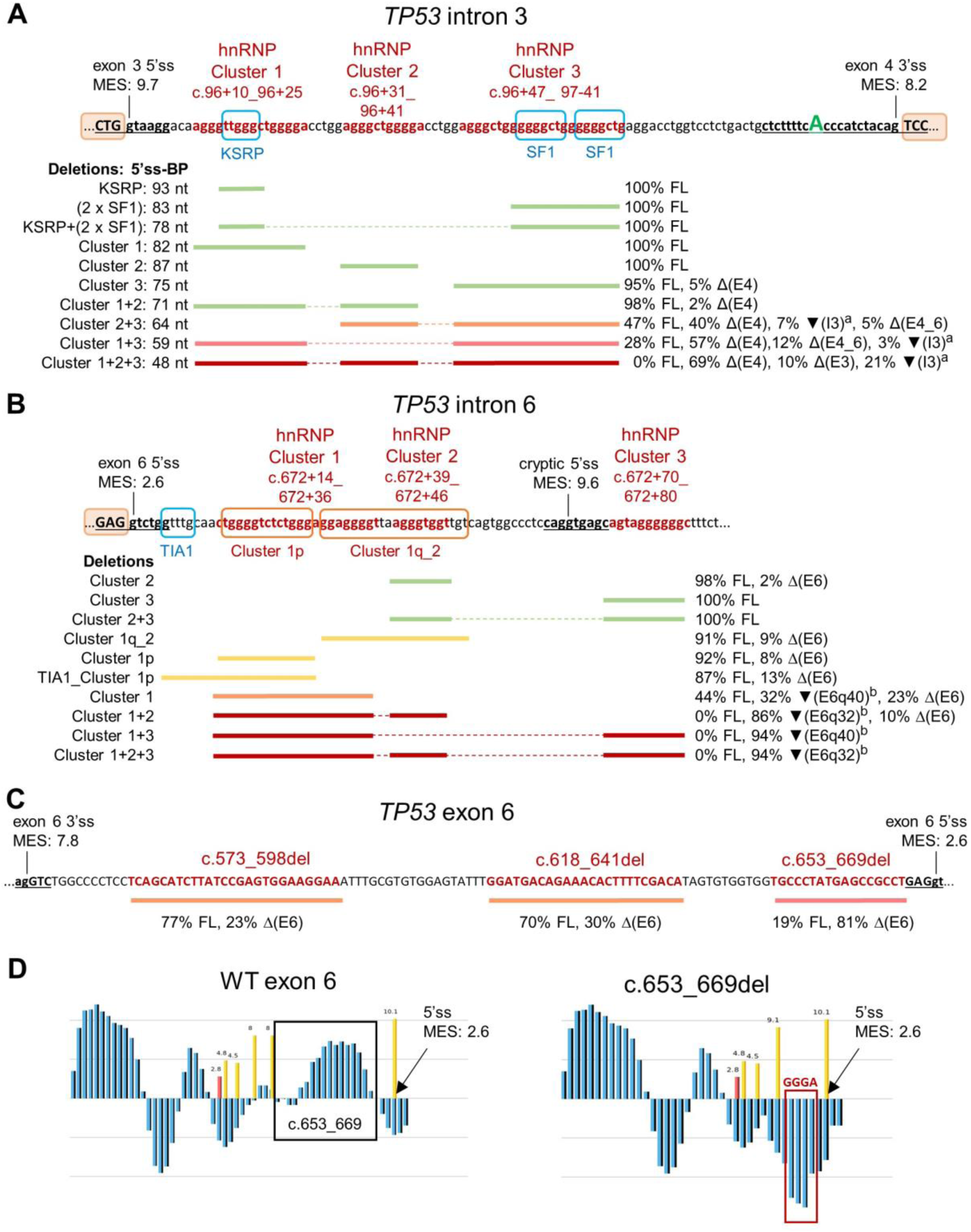
Microdeletion analysis of *TP53* intron 3, intron 6, and exon 6: SRE mapping and assay results. **A)** Intron 3 G-rich intervals and predicted binding sites for splicing factors KSRP, SF1, and hnRNPs A1 or F/H group. The experimentally inferred canonical adenosine BP (c.97-12A)^44^ is capitalized and in green font. ^a^The ▼(I3) transcripts retained the intron 3 sequence with intronic microdeletions. **B)** Intron 6 predicted binding site for TIA1 splicing factor and G-rich intervals that are predicted binding sites for hnRNPs A1 or F/H group. ^b^The ▼(E6q32) and ▼(E6q40) transcripts were a product of intronic cryptic donor usage located at c.672+63 with MES score 9.6. The percentage of uncharacterized transcripts are not shown. **C)** ESE-rich intervals in exon 6 with different levels of splicing impact upon deletion. **D)** HEXplorer profile of sequence upstream of exon 6 donor site (MES score 2.6); c.653_669del removes the ESE cluster in the WT sequence and creates an ESS motif (GGGA) that binds hnRNP F/H. See Supplementary Table S2 for detailed assay results.

#### Intron 3

The individual deletions of hnRNP cluster 1, hnRNP cluster 2, KSRP, and SF1 in intron 3, as well as the combined deletion of KSRP+SF1, had no effect on splicing (**Figure 2A**). It had been previously reported that deletions of the two tracts of six Gs in intron 3 (c.96+53_97-52, c.97-49_97-44) induced intron 2 retention^12^, but we did not observe this event in our splicing assay of the hnRNP cluster 3 microdeletion spanning both of these G-runs (c.96+47_ 97-41del). Instead, c.96+47_ 97-41del induced exon 4 skipping (5% △(E4)).

The combined deletions of intron 3 hnRNP clusters also induced exon 4 skipping (2-69% △(E4)) and other minor aberrations including exon 3 skipping, intron 3 retention, and skipping of exons 4 to 6 (**Figure 2A**). The consequent FL transcript depletion correlated with the donor-to-BP distance resulting from the deletions (**Figure 2A**): clusters 1+2 (98% FL, 71-nt distance); 2+3 (47% FL, 64-nt distance); 1+3 (28% FL, 59-nt distance); and 1+2+3 (0% FL, 48-nt distance). It is likely that the splicing impact of the last three combined deletions is not hnRNP-related, but rather due to spatial constraint by shortening the distance between the exon 3 donor and the experimentally inferred canonical adenosine BP in intron 3 located at c.97-12^44–46^. Notably, the intron 3 cluster 1+2+3 deletion shortened the distance between the donor and the c.97-12A BP from 97 nt to 48 nt. Our observation of complete splicing impact by intron 3 cluster 1+2+3 deletion is in agreement with the findings from a previous study^46^ that established a donor-to-BP length cutoff of <50 nt for critical risk of mis-splicing and profound splicing abnormalities.

To test the relevance of the intron 3 sequence in the splicing process, we replaced it with *ATM* intron 21 sequence, which has a similar size (114 nt) and does not contain G-runs. This chimeric construct only induced a low level of exon 4 skipping (1.3% △(E4), **Supplementary Table S2**), suggesting that splicing factors other than hnRNP F/H may be involved in *TP53* exon 3 and 4 recognition in the SKBR3 cell line. This intron 3 substitution assay did not result in intron 2 retention, also suggesting that G-quadruplex structures may not be necessary for normal splicing of this region. Moreover, the presence of a minor multi-exon skipping transcript lacking exons 4, 5 and 6 in the assay of intron 3 clusters 2+3 and 1+3 deletions (**Figure 2A**) suggests some interdependence between these three exons for their recognition by the splicing machinery.

#### Intron 6

Analysis of intron 6 microdeletions revealed that the number of ISE clusters or G-runs downstream of the native donor influenced the level of weak donor site usage (**Figure 2B**). Deletion of hnRNP cluster 2 only, and cluster 3 only, each containing one G-run, had negligible or no splicing impact (98-100% FL transcript). Deletion of intron 6 hnRNP cluster 1p (c.672+15_672+27del), and cluster 1q_2 (c.672+29_672+49del), each spanning two G-runs, led to minor exon 6 skipping (8-9% △(E6)). Extending the cluster 1p deletion to include the putative TIA1 binding site (c.672+7_672+27del) increased the △(E6) level to 13%. Deletion of cluster 1 (c.672+14_672+36del), spanning three G-runs, induced 23% △(E6) and activation of cryptic donor in intron 6 (32% ▼(E6q40)). Combined deletions of intron 6 hnRNP clusters 1+2, 1+3, and 1+2+3 did not produce any trace of the minigene FL transcript, suggesting that these G-rich sequences are critical for normal splicing of this region. Each of these combined deletions mainly activated the intronic c.672+63 cryptic donor (86-94% of overall expression). Moreover, cluster 1+2 deletion that effectively removed all four G-runs between the native and the cryptic donor resulted in complete inactivation of the native donor site. These results also suggest that the G-rich sequences have a greater contribution to exon 6 donor site recognition compared to the TIA1 binding site.

#### Exon 6

Microdeletion analysis of exon 6 HEXplorer-predicted ESE motifs (**Figure 2C**) showed that the three selected microdeletions (c.573_598del, c.618_641del, and c.653_669del) affected exon 6 recognition (23-81% △(E6)), indicating ESE enrichment within these intervals. Additionally, c.653_669del (81% △(E6)) created a G-run upstream and adjacent to the weak native donor (**Figure 2D**). G-runs are known to act as ESS motifs when located within the exon^6^. Specifically, c.653_669del formed the GGGA motif that is the core binding site for hnRNP F/H^6^. SpliceAid predicted the same splicing factors hnRNP F/H to target the intronic G-runs upstream of the c.672+63 cryptic donor, which may also play a role in inhibiting the usage of this cryptic site.

### Analysis of ClinVar-reported deletions

We selected and assayed deletions reported in ClinVar (**Supplemental Table S3**) that partially overlap with intron 3 hnRNP clusters 2+3 (c.96+31_96+54del) or located within the exon 6 ESE-enriched clusters (c.581_585del, c.628_639del and c.655_670del).

In our intron 3 microdeletion analysis (**Figure 2A**), we showed that the hnRNP clusters 2+3 deletion altered splicing (47% FL), and we postulated that splicing alteration was not caused by deletion of specific hnRNP-targeted sequences, but rather by critical shortening between the exon 3 donor site and the canonical adenosine BP (c.97-12A). The ClinVar-reported c.96+31_96+54del had minimal impact on splicing (95% FL). This deletion spanned intron 3 hnRNP clusters 2 and 3 (the latter, partially), shortening the distance between exon 3 donor and c.97-12A to 73 nt, further supporting the claim that it was actually donor-to-BP distance that explains the splicing abnormality in the microdeletion experiments summarized in **Figure 2A**. The c.96+31_96+54del variant is currently classified as likely benign (ClinVar Accession: VCV000926699.2, accessed 24 September 2024) with no splicing assay data; evidence of minimal splicing impact from this study supports this likely benign classification.

Of the three ClinVar-reported exon 6 deletions, only the c.655_670del variant had a major impact on splicing (**Supplemental Figure S5**), mainly due to the creation of a new donor within exon 6 that produced an in-frame transcript with a 21 nt deletion (90% △(E6q21)). The pathogenic assertion for this variant in ClinVar is based on the assumption that it causes a translational frameshift (ClinVar Accession: VCV000486553.5, accessed 24 September 2024). Here we show that it actually affects function through aberrant splicing mechanism resulting in the removal of seven amino acids in a clinically relevant domain.

### Analysis of single nucleotide variants

We assayed 134 SNVs, including four variants located at the splice donor/acceptor ±1,2 dinucleotide positions as positive controls, 66 SNVs in exon 3, 43 SNVs in exon 6, and 21 SNVs in intron 6 (**Supplementary Table S3**). Due to the exploratory nature of this study, we assayed all possible SNVs in exon 3 (from c.75 to c.96 totaling 66 variants) to check for exonic variants that affect the recognition of this microexon. We selected exon 6 SNVs with HEXplorer delta HZEI score ≤ -5 and located within the three microdeletions confirmed to result in exon skipping. For intron 6, we selected SNVs between the native and cryptic donors and outside the consensus motifs (c.672+7 to c.672+60) that passed at least two of the following filters: HEXplorer delta HZEI ≥5, PhyloP conservation score >0.5, or SpliceAI max delta score >0.1. Expectedly, 15 of the 21 intronic SNVs that passed these criteria are located within G-runs. We excluded exon 6 and intron 6 SNVs with gnomAD minor allele frequency >0.0001. SNVs that impact splicing (<95% FL transcript) are shown in **Table 1**.

**Table 1.** Percentage of transcripts produced by spliceogenic *TP53* SNVs.

| Variant | Exon (E)/ Intron (I) | Full-length Transcript | PTC transcripts | In-frame and other transcripts |
| --- | --- | --- | --- | --- |
| c.75-2A>G | I2 | - | 28% $\Delta$(E3)<br>34% $\Delta$(E3_E4p19) | 31% $\Delta$(E3_E4)<br>4% $\nabla$(I2)<br>3% 785-nt uncharacterized transcript |
| c.82G>T | E3 | 85% | 5% $\Delta$(E6)<br>10% $\Delta$(E3) | - |
| c.96G>T | E3 | 83% | 17% $\Delta$(E3) | - |
| c.96+1G>T | I3 | - | 100% $\Delta$(E3) | - |
| c.560-2A>C | I5 | - | 78% $\Delta$(E6p17)<br>17% $\Delta$(E6) | 3% $\nabla$(I5)<br>3% 983-nt uncharacterized transcript |
| c.592G>T | E6 | 74% | 26% $\Delta$(E6) | - |
| c.593A>T | E6 | 80% | 20% $\Delta$(E6) | - |
| c.619G>T | E6 | 91% | 9% $\Delta$(E6) | - |
| c.622G>T | E6 | 84% | 16% $\Delta$(E6) | - |
| c.661G>T | E6 | 92% | 8% $\Delta$(E6) | - |
| c.662A>G | E6 | 92% | 8% $\Delta$(E6) | - |
| c.662A>T | E6 | 90% | 10% $\Delta$(E6) | - |
| c.672+1G>A | I6 | - | 90% $\nabla$(E6q5)<br>6% $\Delta$(E6) | 2% 925-nt, 2% 1005-nt uncharacterized transcripts |
| c.672+7T>C | I6 | 89% | 11% $\Delta$(E6) | - |
| c.672+7T>G | I6 | 90% | 10% $\Delta$(E6) | - |
| c.672+18G>A | I6 | 86% | 14% $\Delta$(E6) | - |
| c.672+26G>C | I6 | 92% | 8% $\Delta$(E6) | - |

#### Control variants

We assayed four splice site dinucleotide variants flanking exon 3 (c.75-2A>G, c.96+1G>T) and exon 6 (c.560-2A>C, c.672+1G>A), assumed from position and bioinformatic prediction to impact splicing, as positive controls. The four variants totally disrupted splicing without any trace of the minigene FL transcript, generating at least 11 anomalous transcripts: △(E3), △(E3_E4p19), △(E3_E4), ▼(I2), △(E6p17), △(E6), ▼(I5) and ▼(E6q5), as well as three minor uncharacterized transcripts of 785, 925 and 1005 nucleotides. Only c.96+1G>T induced complete exon skipping (100% △(E3)) while the remaining variants induced combinations of aberrant transcripts with alternative splice site usage, (multi-)exon skipping, or intron retention. The exon 6 acceptor site variant c.560-2A>C principally generated △(E6p17) transcript (78%), resulting from the use of an exonic cryptic acceptor site 17 nt downstream strengthened by this variant (MES: 2.58 → 4.30). The exon 6 donor site variant c.672+1G>A induced 90% ▼(E6q5) transcript due to the usage of a weak cryptic donor (MES 0.72) 5 nt downstream, whose recognition is likely promoted by the surrounding SREs. This ▼(E6q5) transcript was also previously detected in the KG-1 human myeloid leukemia cell line that harbors the c.672+1G>A variant^47^. However, RT-PCR analysis of mRNA (without NMD inhibition) from peripheral blood mononuclear cells of a patient with Li–Fraumeni-like syndrome showed that germline c.672+1G>A induced an in-frame ▼(E6q18) transcript^48^. This 18-nt partial intron retention was caused by the activation of another intronic cryptic donor that overlaps with the first G-run in intron 6 hnRNP cluster 1.

#### Test variants

Of the exon 3 SNVs located outside of the acceptor and donor consensus motifs, only the c.82G>T variant resulted in exon 3 skipping (10% △(E3)); this variant also had a minor effect on exon 6 recognition (5% △(E6)). The remaining exon 3 SNVs had no impact on splicing, except c.96G>T (17% △(E3)), which is the last base of the exon and part of the donor consensus motif. Seven exon 6 SNVs (c.592G>T, c.593A>T, c.619G>T, c.622G>T, c.661G>T, c.662A>G, and c.662A>T) and four intron 6 SNVs (c.672+7T>C, c.672+7T>G, c.672+18G>A, and c.672+26G>C) disrupted exon 6 recognition (8-26% △(E6)). The remaining exon 6 and intron 6 SNVs resulted in either very low level △(E6) transcript or 100% FL transcript.

#### DeepCLIP analysis

In order to identify the putative splicing factors involved in exon 3 and exon 6 recognition, and to determine the effect of SNVs on the binding of these factors, we ran DeepCLIP predictions for 12 SNVs outside of consensus splice site motifs that resulted in exon skipping (**Table 2, Supplementary Table S4**). The delta HZEI scores generally correlated with DeepCLIP predictions. For example, delta HZEI scores indicating enhancer motif loss had corresponding DeepCLIP-predicted decrease in the binding of enhancer proteins.

**Table 2.**
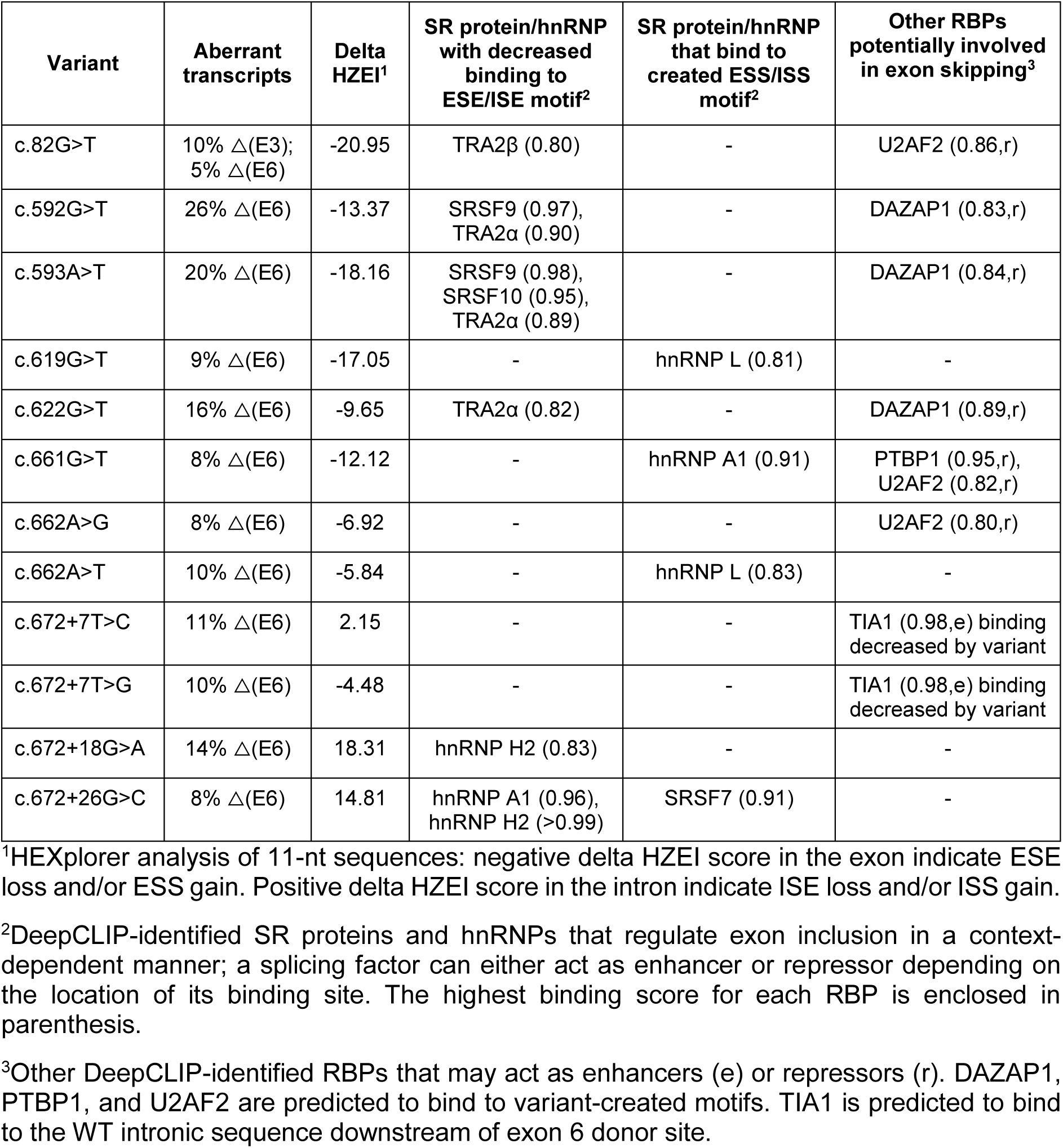
HEXplorer delta HZEI and DeepCLIP analysis of SNVs outside the consensus splice site motifs and confirmed to result in exon skipping.

| Variant | Aberrant transcripts | Delta HZEI<sup>1</sup> | SR protein/hnRNP with decreased binding to ESE/ISE motif<sup>2</sup> | SR protein/hnRNP that bind to created ESS/ISS motif<sup>2</sup> | Other RBPs potentially involved in exon skipping<sup>3</sup> |
| --- | --- | --- | --- | --- | --- |
| c.82G>T | 10% $\Delta$(E3); 5% $\Delta$(E6) | -20.95 | TRA2β (0.80) | - | U2AF2 (0.86,r) |
| c.592G>T | 26% $\Delta$(E6) | -13.37 | SRSF9 (0.97), TRA2α (0.90) | - | DAZAP1 (0.83,r) |
| c.593A>T | 20% $\Delta$(E6) | -18.16 | SRSF9 (0.98), SRSF10 (0.95), TRA2α (0.89) | - | DAZAP1 (0.84,r) |
| c.619G>T | 9% $\Delta$(E6) | -17.05 | - | hnRNP L (0.81) | - |
| c.622G>T | 16% $\Delta$(E6) | -9.65 | TRA2α (0.82) | - | DAZAP1 (0.89,r) |
| c.661G>T | 8% $\Delta$(E6) | -12.12 | - | hnRNP A1 (0.91) | PTBP1 (0.95,r), U2AF2 (0.82,r) |
| c.662A>G | 8% $\Delta$(E6) | -6.92 | - | - | U2AF2 (0.80,r) |
| c.662A>T | 10% $\Delta$(E6) | -5.84 | - | hnRNP L (0.83) | - |
| c.672+7T>C | 11% $\Delta$(E6) | 2.15 | - | - | TIA1 (0.98,e) binding decreased by variant |
| c.672+7T>G | 10% $\Delta$(E6) | -4.48 | - | - | TIA1 (0.98,e) binding decreased by variant |
| c.672+18G>A | 14% $\Delta$(E6) | 18.31 | hnRNP H2 (0.83) | - | - |
| c.672+26G>C | 8% $\Delta$(E6) | 14.81 | hnRNP A1 (0.96), hnRNP H2 (>0.99) | SRSF7 (0.91) | - |

<sup>1</sup>HEXplorer analysis of 11-nt sequences: negative delta HZEI score in the exon indicate ESE loss and/or ESS gain. Positive delta HZEI score in the intron indicate ISE loss and/or ISS gain.
<sup>2</sup>DeepCLIP-identified SR proteins and hnRNPs that regulate exon inclusion in a context-dependent manner; a splicing factor can either act as enhancer or repressor depending on the location of its binding site. The highest binding score for each RBP is enclosed in parenthesis.
<sup>3</sup>Other DeepCLIP-identified RBPs that may act as enhancers (e) or repressors (r). DAZAP1, PTBP1, and U2AF2 are predicted to bind to variant-created motifs. TIA1 is predicted to bind to the WT intronic sequence downstream of exon 6 donor site.

DeepCLIP analysis identified at least seven SR proteins and hnRNPs that bind to SREs in WT *TP53* pre-mRNA (SRSF9, SRSF10, TRA2α, TRA2β, hnRNP A1, and hnRNP H2), ensuring normal splicing of exons 3 and 6. Six of the spliceogenic variants disrupted the ESE/ISE motifs targeted by these enhancer proteins. ESE disruption by exon 3 variant c.82G>T decreased TRA2β binding resulting in decreased exon 3 inclusion. Similarly, exon 6 variants c.592G>T, c.593A>T, and c.622G>T weakened the ESEs leading to decreased binding of SRSF9, SRSF10, or TRA2α. Intron 6 variants c.672+18G>A and c.672+26G>C disrupted the ISE motifs/G-runs resulting in decreased binding of hnRNPs A1 or H2. Predicted decreased binding of these enhancer proteins in exon 6 or intron 6 could explain the decreased exon 6 inclusion.

Five of the six variants that disrupted enhancer motifs (c.82G>T, c.592G>T, c.593A>T, c.622G>T, and c.672+26G>C) also created binding sites for repressor proteins U2AF2, PTBP1, DAZAP1, or SRSF7. Ectopic binding of PPT splicing factors U2AF2 and PTBP1 in the exon can repress exon inclusion^49,50^. Binding of DAZAP1 to the exon^51^ and binding of SRSF7 downstream of the donor splice site^52^ can also inhibit exon inclusion. Moreover, we predicted four exonic variants (c.619G>T, c.661G>T, c.662A>G and c.662A>T) to alter splicing through ESS creation only, which then become target sites for U2AF2, PTBP1, hnRNP A1 or hnRNP L that would function as repressors in this context^49,50,53,54^.

In addition, we predicted the TIA1 protein to bind to the intronic sequence immediately downstream of the weak exon 6 donor site. TIA1 promotes the recognition of weak donor sites when bound to the downstream intronic uridine-rich sequences by stabilizing U1 snRNP recruitment^43^. The decrease in exon 6 inclusion resulting from c.672+7T>C and c.672+7T>G variants could be due to TIA1 binding site disruption. Notably, this putative TIA1 binding site is also a weak cryptic donor motif (MES 0.72), the one that generates the ▼(E6q5) transcript when activated. We hypothesize that TIA1 binding to this uridine-rich motif prevents the usage of this cryptic donor while simultaneously stabilizing U1 snRNP binding to the adjacent weak native donor site. In the event of native donor site abrogation, as in the case of c.672+1G>A (**Table 1**), this nearby cryptic donor would then be used. We note that the TIA1 binding site alteration had a modest effect, from 8% when disrupted by SNVs (**Table 2**) to 13% when deleted (**Figure 2B**), so it is likely that TIA1 is one more splicing factor of the set involved in splicing control of this region.

### Prediction of splicing impact: SpliceAI vs delta HZEI

We obtained SpliceAI predictions for all individual deletions and SNVs assayed (**Supplementary Table S5, Figure 3A**). Expectedly, the four SNVs located at the donor/acceptor ±1,2 dinucleotide positions with complete splicing impact had the highest SpliceAI scores (0.98-1.0).

**Figure 3.**
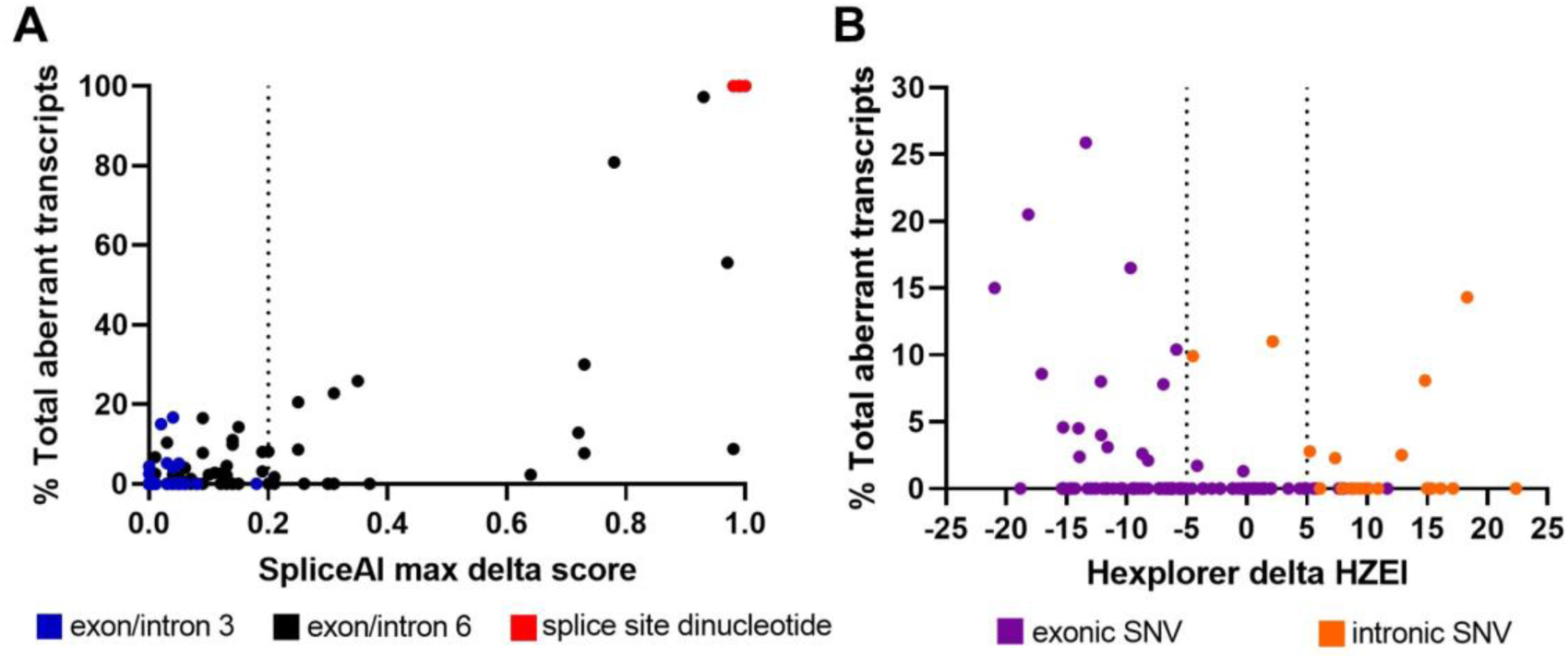
Splicing impact prediction. **A)** SpliceAI scores and percentage of total aberrant transcripts for all individual deletions and SNVs assayed (n=152). Variants with SpliceAI score ≥0.2 are predicted to alter splicing^36^. **B)** HEXplorer delta HZEI scores and percentage of total aberrant transcripts for SNVs outside of splice consensus motifs (n=112). Exonic SNVs with delta HZEI score ≤-5 and intronic SNVs with delta HZEI score ≥5 are predicted to disrupt SREs^34^.

We computed the likelihood ratio for spliceogenicity using the SpliceAI thresholds previously recommended for bioinformatic prediction of splicing impact for variants outside of the donor/acceptor ±1,2 dinucleotide positions, based on analysis of experimental data without quantification of transcript levels^36^. Likelihood ratio analysis based on quantitative data from this study, and selecting ≥5% aberrant transcript as a splice event (**Table 3**), generated similar findings to those reported previously^36^. The SpliceAI cutoff score of ≥0.2 equated to a moderate level of evidence for spliceogenicity. Of the 20 variants with SpliceAI score ≥0.2, 12 were true positive calls. Seven of these true positive calls, including five deletions (c.573_598del, c.618_641del, c.653_669del, c.655_670del, and c.672+14_672+36del) and two SNVs (c.592G>T and c.593A>T), led to >20% expression of aberrant transcripts. Regarding the mechanism of splicing impact of the 12 spliceogenic variants with SpliceAI score ≥0.2, seven were ESE/ISE motif deletions, four were SRE disruption by SNVs, and one was a deletion that created a new donor site. The remaining eight variants with SpliceAI score ≥0.2 were false positives and had aberrant transcript levels ranging from 0-2.3%. SpliceAI score >0.1 and <0.2 equated to uninformative strength of evidence, and the level of aberrant transcripts ranged from 0-14.3% for the 22 variants in this category. The SpliceAI cutoff score of ≤0.1 equated to a supporting level of evidence against spliceogenicity. It is notable that the eight variants in this category that were observed to impact splicing (i.e. false negative calls) had only 5.1-16.7% expression of aberrant transcripts. These eight false negatives were comprised of four SRE-disrupting SNVs, one deletion within an ESE cluster in exon 6, one exon 3 SNV within the donor consensus motif, and two intron 3 deletions that shortened the intron size. Interestingly, all exon 3 SNVs and intron 3 individual deletions had SpliceAI score <0.2, in general agreement with the assay results, suggesting a sequence context that ensures normal splicing or only minimal (≤16.7%) splicing aberration in this region.

**Table 3.** Likelihood ratio analysis of the maximum SpliceAI delta score for *TP53* deletions and SNVs outside the donor/acceptor ±1,2 dinucleotide positions using recommended thresholds.

| SpliceAI delta score | Splice event - no (<5% aberrant transcript) | | Splice event - yes ($\geq 5\%$ aberrant transcript) | | LR | Low CI | High CI | Evidence strength |
| --- | --- | --- | --- | --- | --- | --- | --- | --- |
|  | n | Proportion | n | Proportion |  |  |  |  |
| $\leq 0.1$ | 98 | 0.79 | 8 | 0.33 | 0.42 | 0.24 | 0.75 | Supporting evidence (non-spliceogenicity) |
| $> 0.1$ and $< 0.2$ | 18 | 0.15 | 4 | 0.17 | 1.15 | 0.43 | 3.09 | Uninformative |
| $\geq 0.2$ | 8 | 0.06 | 12 | 0.50 | 7.75 | 3.55 | 16.92 | Moderate evidence (spliceogenicity) |

Abbreviation: CI, confidence interval; LR, likelihood ratio; n, number.

To compare the performance of bioinformatic tools in predicting spliceogenic SNVs that act through SRE-disruption, we selected 112 SNVs (12 spliceogenic and 100 non-spliceogenic) located outside of the consensus motifs (**Supplementary Table S6**). SpliceAI had a lower sensitivity (33%) than the HEXplorer 11-nt delta HZEI (83%) for spliceogenic SNVs that disrupt exonic or intronic SREs. However, SpliceAI had a higher specificity (76%) than the 11-nt delta HZEI (37%). While the delta HZEI score is useful in determining variants that cause ESE/ISE loss and/or ESS/ISS gain, this tool performed poorly at predicting SRE-disrupting variants that actually alter splicing. The delta HZEI cutoff score ≤ -5 for exon and ≥5 for intron had a high false positive rate of 63% for exonic and intronic variants outside of consensus splice motifs (**Figure 3B, Supplementary Table S6**). For exonic SNVs only (8 spliceogenic and 83 non-spliceogenic), we observed that a different approach using delta HZEI cutoff score of ≤ -40 for whole exons would lower the false positive rate (35%) compared to the 11-nt delta HZEI (55%). The sensitivity of bioinformatic tools for splicing impact of exonic SRE-disrupting SNVs were: 100% for 11-nt delta HZEI, 88% for whole exon delta HZEI, and 38% for SpliceAI. Inversely, specificity for exonic SRE-disrupting SNVs were 80% for SpliceAI, 65% for whole exon delta HZEI, and 45% for 11-nt delta HZEI. Overall, SpliceAI and HEXplorer had limitations when predicting the splicing impact of variants acting via SRE mechanism.

## Discussion

We constructed the mgTP53_2-9 minigene containing *TP53* exons 2 to 9, a construct that can be used for splicing studies of any variant or motif within the inserted *TP53* region. This include variations that induce mRNA transcripts encoding truncated p53 isoforms, cause in-frame deletions, or generate transcripts sensitive to NMD. In this study, we used this minigene construct to examine the role of SREs on microexon 3 and exon 6 recognition. This is the first study that analyzed the effect of *TP53* deletions and SNVs on SRE motifs and the predicted binding of corresponding splicing factors. Our minigene assays that measured the splicing impact of deletions and SNVs within SRE-rich regions has provided mechanistic insights on normal and impaired splicing of the selected exons. In addition, we verified that adequate donor-to-BP distance is required for normal splicing.

Exon 3, which has very strong acceptor (MES 11.65) and donor (MES 9.68) splice sites, was tolerant to disruption of SRE motifs within the exon and the downstream intron 3. Our minigene assays in SKBR3 cells showed that deletion of G-quadruplex structures in intron 3 did not induce intron 2 retention. Further, replacement of the entire intron 3 sequence with *ATM* intron 21 that is devoid of G-runs did not affect intron 2 excision in SKBR3 cells, supporting the notion that normal splicing of this region proceeds even in the absence of G-quadruplexes. These findings contrast previous results of intron 3 G-run deletion and mutagenesis experiments using a green fluorescent protein reporter containing *TP53* exons 2 to 4 assayed in H1299 cells^12^. This discordance in observations of intron 2 retention could possibly be ascribed to differences in cell lines and minigene construct design.

On the other hand, our findings showed the importance of intronic G-runs in exon 6 splicing, particularly in the activity of the weak exon 6 donor site (MES 2.6). There are five G-runs within the 85-nt intronic sequence directly downstream of exon 6. Deletion of three G-runs (hnRNP cluster 1) decreased the percentage of FL transcript by more than half; deletion of at least four G-runs (hnRNP clusters 1+2 or 1+3) was enough to completely inactivate the exon 6 donor site and activate the strong intronic cryptic donor site (MES 9.6). ESE motifs in exon 6 also contribute to normal exon 6 splicing, and ESE motif deletions resulted in decreased exon 6 inclusion. Overall, we demonstrated that *TP53* deletions spanning multiple enhancer binding sites and located close to the regulated splice sites led to drastic splicing alterations. Whereas, SRE-disrupting SNVs induced weak splicing effects due to SRE motif redundancy, motif location, sequence context, or cooperation of several different splicing motifs and RBPs for exon recognition, similar to results of previous studies^55–57^. Therefore, in the context of germline variant classification, SRE-disrupting SNVs are less likely to confer pathogenicity via severe impact on mRNA processing. Nevertheless, there have been described SRE-disrupting SNVs that induce total splicing impact (no FL or negligible amounts), such as *CHEK2* c.883G>A, c.883G>T and c.884A>T that were classified as pathogenic/likely pathogenic variants^58^.

SpliceAI, a deep neural network considered as one of the best splicing prediction tools to date^59^, positively detected all SRE-disrupting deletions and SNVs located outside of the splice consensus motifs that produced >20% aberrant transcripts, using the threshold of ≥0.2. This SpliceAI threshold was recommended conservatively for the application of supporting level of evidence for predicted impact on splicing within the ACMG/AMP framework^36^. Notably, likelihood ratio estimation derived from this dataset, enriched for SNVs and deletions affecting SRE motifs, provided further justification for SpliceAI score ≥0.2 as a conservative cutoff for the application of supporting level of bioinformatic evidence. SpliceAI that captures sequence features up to 10,000 nt surrounding a variant^35^, had a lower false positive rate than the SRE-specific tool HEXplorer delta HZEI, that only interrogates short stretches of nucleotide sequence. HEXplorer’s high false positive rate was probably due to SRE motif location and redundancy. Disruption of one enhancer motif while other enhancer motifs remain intact is less likely to cause a splicing aberration, especially if the motif is distantly located from the regulated splice site. However, SRE- and RBP-specific tools (HEXplorer, SpliceAid and DeepCLIP) are useful in mapping *cis*-regulatory elements and in unravelling the probable molecular mechanism and splicing factors underlying an experimentally observed splicing impact. Irrespective of the prediction tool selected, existence of both false positive and false negative predictions means that experimental assessment of variant impact on splicing remains important for variant classification.

We have identified the splicing factors TRA2β in exon 3; SRSF9, SRSF10, and TRA2α in exon 6; and hnRNP A1, hnRNP H2, and TIA1 in intron 6 as putative enhancers involved in *TP53* pre-mRNA splicing regulation. However, TRA2β may only have a minor role in exon 3 recognition as this exon is bounded by very strong acceptor and donor splice site motifs. We also identified putative repressor proteins that bind to variant-created silencer motifs including hnRNP A1, hnRNP L, SRSF7, DAZAP1, PTBP1, and U2AF2. It is possible that other splicing factors may be involved in regulating *TP53* splicing via SRE binding; we provide an extended list of DeepCLIP-identified splicing factors for the selected *TP53* regions in **Supplementary Table S4**.

We emphasize the role of predicted splicing factors targeting intron 6 elements to promote exon 6 donor site activity. Namely, TIA1 that binds directly downstream of the weak donor site, and hnRNP A1 and hnRNP H that bind to G-runs. TIA1, which is also a tumor suppressor^60^, has been implicated in p53 mRNA translational control in B cells by binding to the 3’ untranslated region^61^. A previous study showed that hnRNP F/H interacts with a G-quadruplex structure located at the *TP53* pre-mRNA polyadenylation signal in DNA-damaged cells; consequently, hnRNP F/H depletion compromises pre-mRNA 3′-end processing, p53 expression, and p53-mediated apoptosis^62^. Here, we predict that TIA1 and hnRNP H are also potentially involved in normal exon 6 splicing, preventing exon 6 skipping and/or intron 6 cryptic donor activation that would generate mRNA transcripts with a premature termination codon. It would be interesting to investigate the effect of depletion of TIA1, hnRNP A1, and hnRNP H, as well as the other splicing factors mentioned above, on *TP53* pre-mRNA splicing. It was previously suggested that deregulation of proper splicing factor balance in tumors, in addition to genetic variants affecting the *cis*-regulatory elements, could activate the intron 6 cryptic acceptor site leading to the expression of prometastatic p53Ψ isoform^14^. Future studies may also look into the potential production of a C-terminal truncated p53 isoform resulting from intron 6 cryptic donor activation upon splicing factor depletion (e.g. TIA1, hnRNP A1, and hnRNP H) or *cis*-regulatory element deletion/variation, and the functional role of such an isoform. It is worth noting that we used the SKBR3 breast cancer cell line in our assays; the splicing profile resulting from variations may be different in other cell lines as different splicing factors may also be involved.

Outside of SRE studies, we confirmed findings from a previous minigene study of *HBB* intron 1^46^ that a donor-to-BP distance of <50 nt can cause completely abnormal splicing. This previous study also established a donor-to-BP distance cutoff of >60 nt for low risk of mis-splicing^46^. However, in our assay of *TP53* intron 3 cluster 2+3 deletion, we found that shortening the donor-to-BP distance to 64 nt still induced aberrant transcripts amounting to 53% of the overall expression. We also found that *TP53* intron 3 deletions with donor-to-BP distance of 71-75 nt resulted in low level exon 4 skipping (2-5%). That is, our findings indicated a longer donor-to-distance cutoff of >75 nt for low risk of abnormal splicing. Future studies using introns from multiple genes as models will further refine the donor-to-BP distance cutoffs for predicting risk for abnormal splicing.

In conclusion, we provided splicing data for 27 deletions, one intron substitution, and 134 SNVs in the *TP53* gene, and we explored and discussed splicing mechanisms to explain the observed splicing impact induced by variants. We demonstrated that intron 6 G-runs hugely contribute to exon 6 splicing regulation. We also provided more data to inform prediction of splicing impact due to intronic deletions that shorten intron size. Therefore, intronic deletions outside the consensus splice motifs can have severe consequences and should be considered for splicing assays irrespective of bioinformatic prediction of impact especially if supported by clinical data.

## URL of databases and online tools

ClinVar (https://www.ncbi.nlm.nih.gov/clinvar/)

DeepCLIP (https://deepclip-web.compbio.sdu.dk/)

HEXplorer (https://rna.hhu.de/HEXplorer/)

gnomAD (https://gnomad.broadinstitute.org/)

Likelihood ratio for spliceogenicity (https://gwiggins.shinyapps.io/lr_shiny/)

MaxEntScan (http://hollywood.mit.edu/burgelab/maxent/Xmaxentscan_scoreseq.html)

SpliceAid (http://www.introni.it/splicing.html)

## Author contributions

**Conceptualization**, DMC; **data curation**, DMC; **formal analysis**, DMC, IL-B, ABS and EAV-S; **funding acquisition**, ABS and EAV-S; **investigation**, IL-B, LS-M, EB-M, and EAV-S; **methodology**, DMC, IL-B, LS-M, EB-M and EAV-S; **supervision**, ABS and EAV-S; **writing— original draft**, DMC, IL-B and EAV-S; **writing—review and editing**, DMC, IL-B, CF, LS-M, EB-M, MdlH, ABS and EAV-S. All authors read and approved the final version of the manuscript.

## Acknowledgements

This work was supported by the Pawlowski Family Gift. EAV-S lab is supported by grants from the Spanish Ministry of Science and Innovation, Acción Estratégica en Salud 2023, ISCIII (PI23/00047) co-funded by FEDER from Regional Development European Funds (European Union). IL-B is supported by a predoctoral from the Consejería de Educación, Junta de Castilla y León (2022–2025). Programa Estratégico Instituto de Biología y Genética Molecular (IBGM), Escalera de Excelencia, Junta de Castilla y León (Ref. CLU-2019-02). ABS and DMC were supported by NHMRC funding (APP177524). The work of CF was supported by a grant from the National Breast Cancer Foundation, Australia (IIRS-21-102). The funders played no role in study design, data collection, analysis and interpretation of data, or the writing of this manuscript. We thank Alicia García-Álvarez for her excellent technical support (EAV-S lab).

## Competing interests

All authors declare no financial or non-financial competing interests.

## Data availability

All data generated or analyzed during this study are included in this published article and its supplementary files.

## Notes

### Competing Interest Statement

The authors have declared no competing interest.

