## Supplementary Figure for "*TP53* minigene analysis of 161 sequence changes provides evidence for role of spatial constraint and regulatory elements on variant-induced splicing impact"

SacI
GAGCTCcttgggttgtggtgaaacattggaagagagaatgtgaagcagccattcttttcctgctccacaggaagccgagctgtct
cagacactggcatggtgttggggagggggttccttctctgcaggcccaggtgacccagggttggaagtgtctcatgctggatccc
cacttttcctcttgcagCAGCCAGACTGCCTTCCGGGTCACTGCCATGGAGGAGCCGCAGTCAGATCCT
AGCGTCGAGCCCCCTCTGAGTCAGGAAACATTTTCAGACCTATGGAAACTgtgagtggatccattggaa Ex2
gggcaggcccaccaccccaccccaaccccagcccctagcagagacctgtgggaagcgaaaattccatgggactgactttctgc
tcttgtctttcagACTTCCTGAAAACAACGTTCTGgtaaggacaagggttgggctggggacctggagggctggggacct Ex3
ggagggctgggggctgggggctgaggacctggtcctctgactgctcttttcacccatctacagTCCCCCTTGCCGTCCCA
AGCAATGGATGATTTGATGCTGTCCCCGGACGATATTGAACAATGGTTCACTGAAGACCCAGGT
CCAGATGAAGCTCCCAGAATGCCAGAGGCTGCTCCCCCCGTGGCCCCTGCACCAGCAGCTCCTAC Ex4
ACCGGCGGCCCCTGCACCAGCCCCCTCCTGGCCCCTGTCATCTTCTGTCCCTTCCCAGAAAACCTA
CCAGGGCAGCTACGGTTTCCGTCTGGGCTTCTTGCATTCTGGGACAGCCAAGTCTGTGACTTGCA
CGgtcagttgccctgaggggctggcttccatgagacttcaatgcctggccgtatcccctgcatttcttttgtttggaactttgggatt
cctcttcaccctttggcttcctgtcagtgtttttttatagtttacccacttaatgtgtgatctctgactcctgtcccaaagttgaatattcc
ccccttgaatttgggcttttatccatcccatcacaccctcagcatctctcctggggatgcagaacttttctttttcttcatccacgtgtat
tccttggcttttgaaaataagctcctgaccaggcttggtggctcacacctgcaatcccagcactctcaaagaggccaaggcaggca
gatcacctgagcccaggagttcaagaccagcctgggtaacatgatgaaacctcgtctctacaaaaaatacaaaaaattagcca
ggcatggtggtgcacacctatagtcccagccacttaggaggctgaggtgggaagatcacttgaggccaggagatggaggctgca
gtgagctgtgatcacaccactgtgctccagcctgagtgacagagcaagaccctatctcaaaaaaaaaaaaaaaaaagaaaagct
cctgaggtgtagacgccaactctctctagctcgctagtgggttgcaggaggtgcttacgcatgtttgtttctttgctgccgtcttccag
ttgctttatctgttcacttgtgccctgactttcaactctgtctccttcctcttcctacagTACTCCCCTGCCCTCAACAAGAT
GTTTTGCCAACTGGCCAAGACCTGCCCTGTGCAGCTGTGGGTTGATTCCACACCCCCGCCCGGCA Ex5
CCCGCGTCCGCGCCATGGCCATCTACAAGCAGTCACAGCACATGACGGAGGTTGTGAGGCGCTG
CCCCCACCATGAGCGCTGCTCAGATAGCGATGgtgagcagctggggctggagagacgacagggctggttgccca
gggtccccaggcctctgattcctcactgattgctcttagGTCTGGCCCCTCCTCAGCATCTTATCCGAGTGGAAGG Ex6
AAATTTGCGTGTGGAGTATTTGGATGACAGAAACACTTTTCGACATAGTGTGGTGGTGCCCTAT
GAGCCGCCTGAGgtctggtttgcaactggggtctctgggaggaggggttaagggtggttgtcagtggccctccaggtgagca
gtagggggctttctcctgctgcttatttgacctccctataaccccatgagatgtgcaaagtaaatgggtttaactattgcacagttg
aaaaaactgaagcttacagaggctaagggcctcccctgcttggctgggcgcagtggctcatgcctgtaatcccagcactttgggag
gccaaggcaggcggatcacgaggttgggagatcgagaccatcctggctaacggtgaaaccccgtctctactgaaaaatacaaaa
aaaaattagccgggcgtggtgctgggcacctgtagtcccagctactcgggaggctgaggaaggagaatggcgtgaacctgggcg
gtggagcttgcagtgagctgagatcacgccactgcactccagcctgggcgacagagcgagattccatctcaaaaaaaaaaaaa
aaggcctcccctgcttgccacaggtctccccaaggcgcactggcctcatcttgggcctgtgttatctcctagGTTGGCTCTGAC Ex7
TGTACCACCATCCACTACAACTACATGTGTAACAGTTCCTGCATGGGCGGCATGAACCGGAGGC
CCATCCTCACCATCATCACACTGGAAGACTCCAGgtcaggagccacttgccaccctgcacactggcctgctgtgcc
ccagcctctgcttgcctctgacccctgggcccacctcttaccgatttcttccatactactacccatccacctctcatcacatccccggc
ggggaatctccttactgctcccactcagttttcttttctctggctttgggacctcttaacctgtggcttctcctccacctacctggagctg
gagcttaggctccagaaaggacaagggtggttgggagtagatggagcctggtttttttaaatgggacaggtaggacctgatttcctt
actgcctcttgcttctcttttcctatcctgagtagTGGTAATCTACTGGGACGGAACAGCTTTGAGGTGCGTGTT Ex8
TGTGCCTGTCCTGGGAGAGACCGGCGCACAGAGGAAGAGAATCTCCGCAAGAAAGGGGAGCCT
CACCACGAGCTGCCCCCAGGGAGCACTAAGCGAGgtaagcaagcaggacaagaagcggtggaggagaccaa
gggtgcagttatgcctcagattcacttttatcacctttccttgcctctttcctagCACTGCCCAACAACACCAGCTCCTCT Ex9
CCCCAGCCAAAGAAGAAACCACTGGATGGAGAATATTTCACCCTTCAGgtactaagtcttggacctctta
tcaagtggaaagtttccagtctaacactcaaaatgccgttttcttcttgactgttttacctgcaattggggcatttgccatcaggggc
agtgatgcctcaaagacaatggctcctggttgtagctaactaacttcagaacaccaacttataccataatatatattttaaaggacc
agaccagctttcaaaaagGAATTC
EcoRI

**Supplementary Figure S1**. **Insert sequence of minigene mgTP53_2-9.** Exons 2 to 9 are shown in orange and upper case, cloning sites (SacI/EcoR1) are indicated in green and underlined and *Alu* sequences are yellow-shadowed. Size of the insert: 3487 bp. Structure: SacI - ivs1 (182 bp) – ex2 (102 bp) – ivs2 (117 bp) – ex3 (22 bp) – ivs3 (109 bp) – ex4 (279 bp) – ivs4 (757 bp) – ex5 (184 bp) – ivs5 (81 bp) – ex6 (113 bp) – ivs6 (568 bp) – ex7 (110 bp) – ivs7 (343 bp) – ex8 (137 bp) – ivs8 (92 bp) – ex9 (74 bp) – ivs9 (217 bp) – EcoR1.

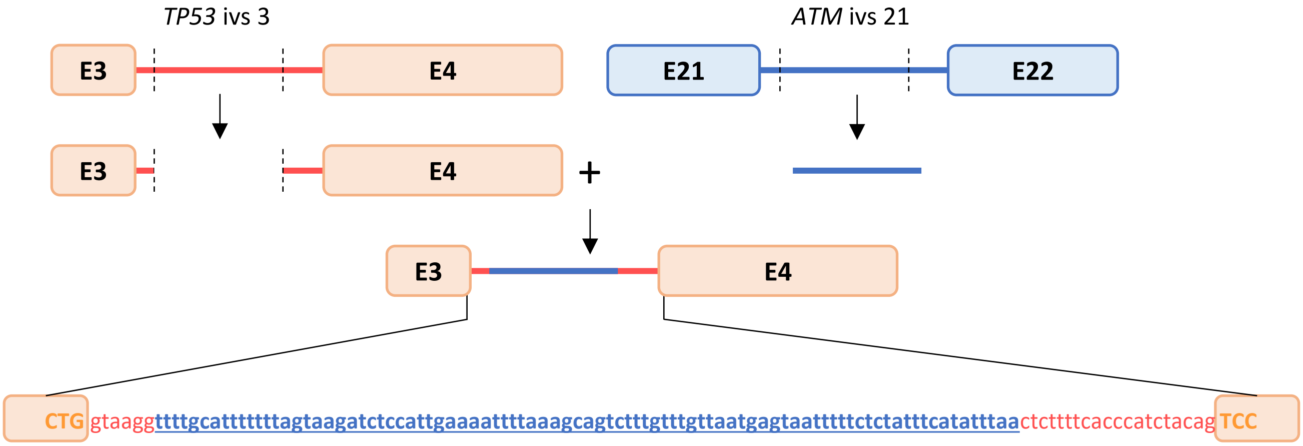


**Supplementary Figure S2**. **Replacement of *TP53* intron 3 with *ATM* intron 21.** Schematic representation of substitution of *TP53* intron 3 by *ATM* intron 21 (shown in blue), maintaining the canonical splice sites of *TP53* exons 3 and 4. The rest of the insert sequence of the mgTP53_2-9 is the same as the one shown in the Supplementary Figure S1.

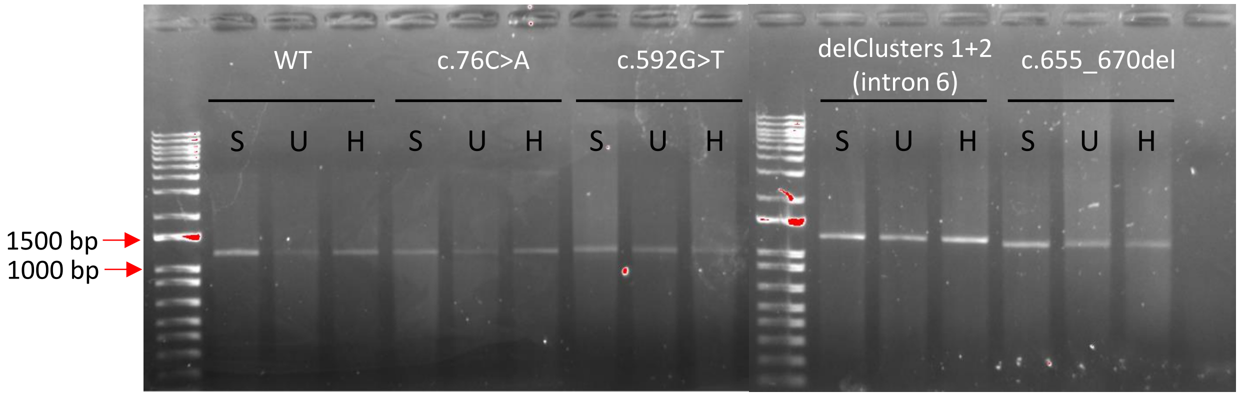


**Supplementary Figure S3. Splicing profile reproducibility in SKBR3, U2OS, and HeLa cell lines.** Agarose gel image showing splicing profiles generated by WT minigene mgTP53_2-9 and four variant constructs: c.76C>A, c.592G>T, c.655_670del, and intron 6 delClusters 1+2 [c.672+14_672+36del;c.672+39_672+46del]. S, SKBR3; U, U2OS; H, HeLa.

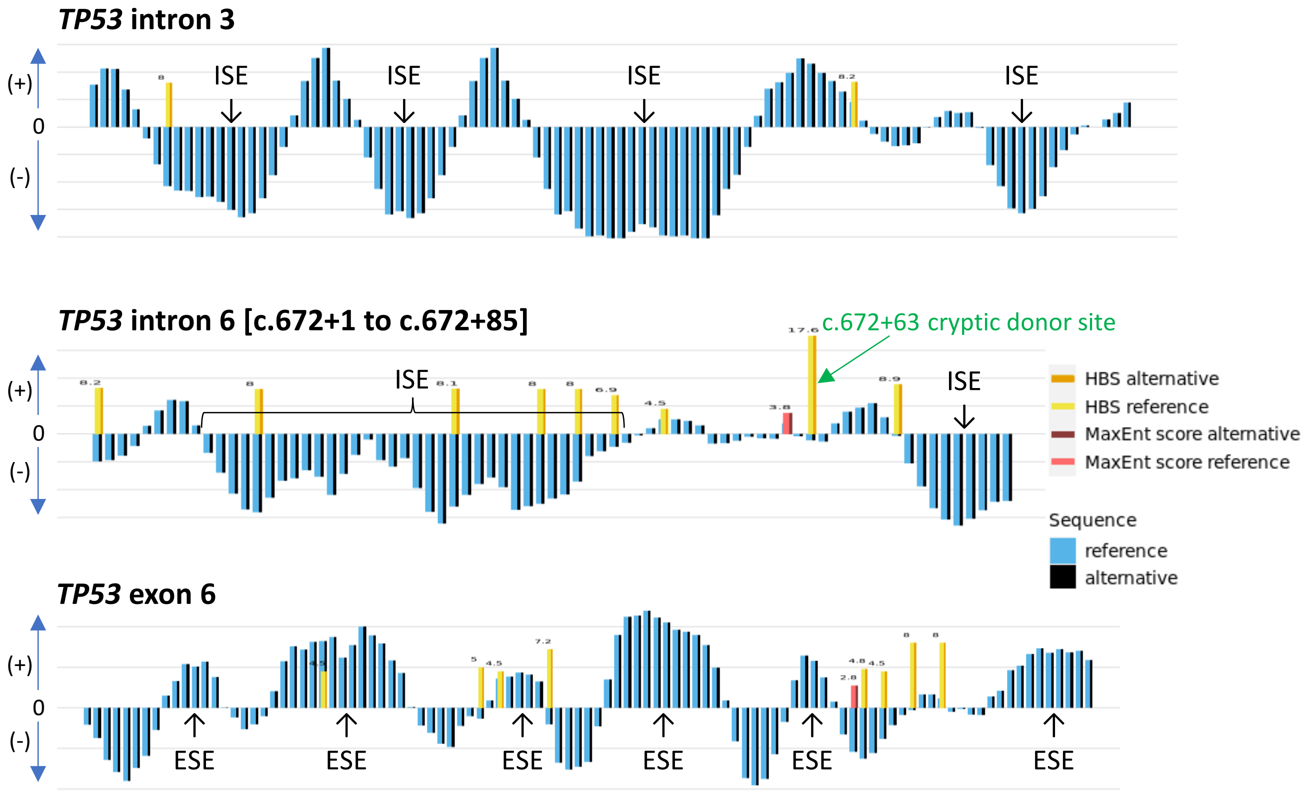


**Supplementary Figure S4**. **HEXplorer profile of *TP53* intron 3, intron 6, and exon 6 WT sequence.** For introns 3 and 6, clusters of negative HEXplorer scores (black/blue bars below zero) represent putative intronic splicing enhancers (ISEs). For exon 6, clusters of positive HEXplorer scores (black/blue bars above zero) represent putative exonic splicing enhancers (ESEs). Cryptic donor (orange/yellow) and acceptor (brown/pink) sites are also shown. None of the cryptic splice sites are predicted as strong motifs except for the c.672+63 cryptic donor site in intron 6.

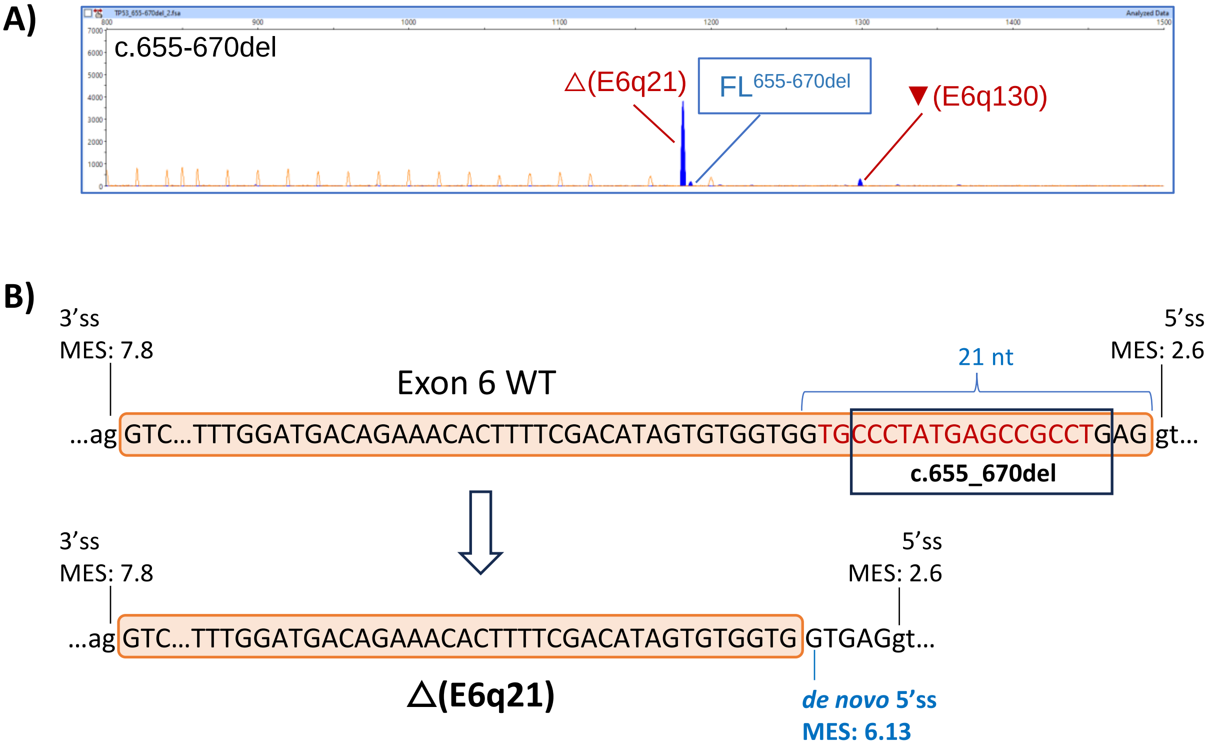


**Supplementary Figure S5**. **Minigene splicing assay result for the ClinVar-reported deletion c.655_670del. A)** Fluorescent fragment analysis result. FAM-labelled products (transcripts, blue peaks; FL$^{655\text{-}670del}$, minigene full-length transcript with the deletion) were run with LIZ1200 (orange peaks) as size standard. The x-axis indicates size in bp and the y-axis represents Relative Fluorescence Units (RFU). **B)** Schematic representation of *de novo* donor site creation within exon 6 owing to c.655_670del (boxed). Usage of the new donor site located 21 nt upstream of the weak exon 6 donor site produced the △(E6q21) in-frame transcript. The ESE-rich sequence affected by c.655_670del is shown in red font.

**A)**

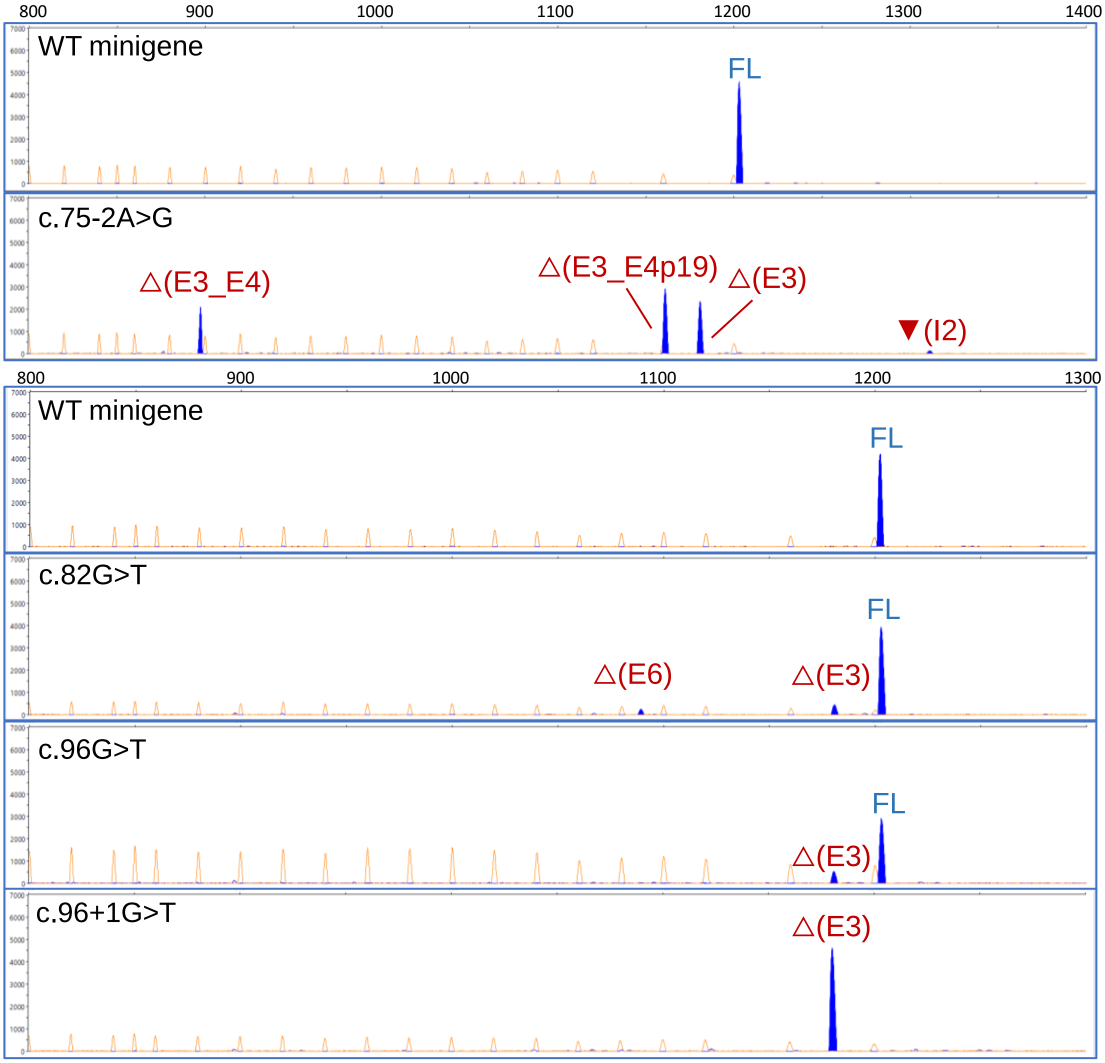


**Supplementary Figure S6. Fluorescent fragment analysis results of spliceogenic single nucleotide variants.** Results for variants that produced <95% full-length transcript and located in: **A)** Exon 3 and flanking splice site ±1,2 dinucleotide positions; **B)** Exon 6 and flanking splice site ±1,2 dinucleotide positions; **C)** Intron 6. FAM-labelled products (transcripts, blue peaks; FL, minigene full-length transcript) were run with LIZ1200 (orange peaks) as size standard. The x-axis indicates size in bp and the y-axis represents Relative Fluorescence Units (RFU).

Note: Supplementary Figure S6 panels B and C are in the next pages.

B)

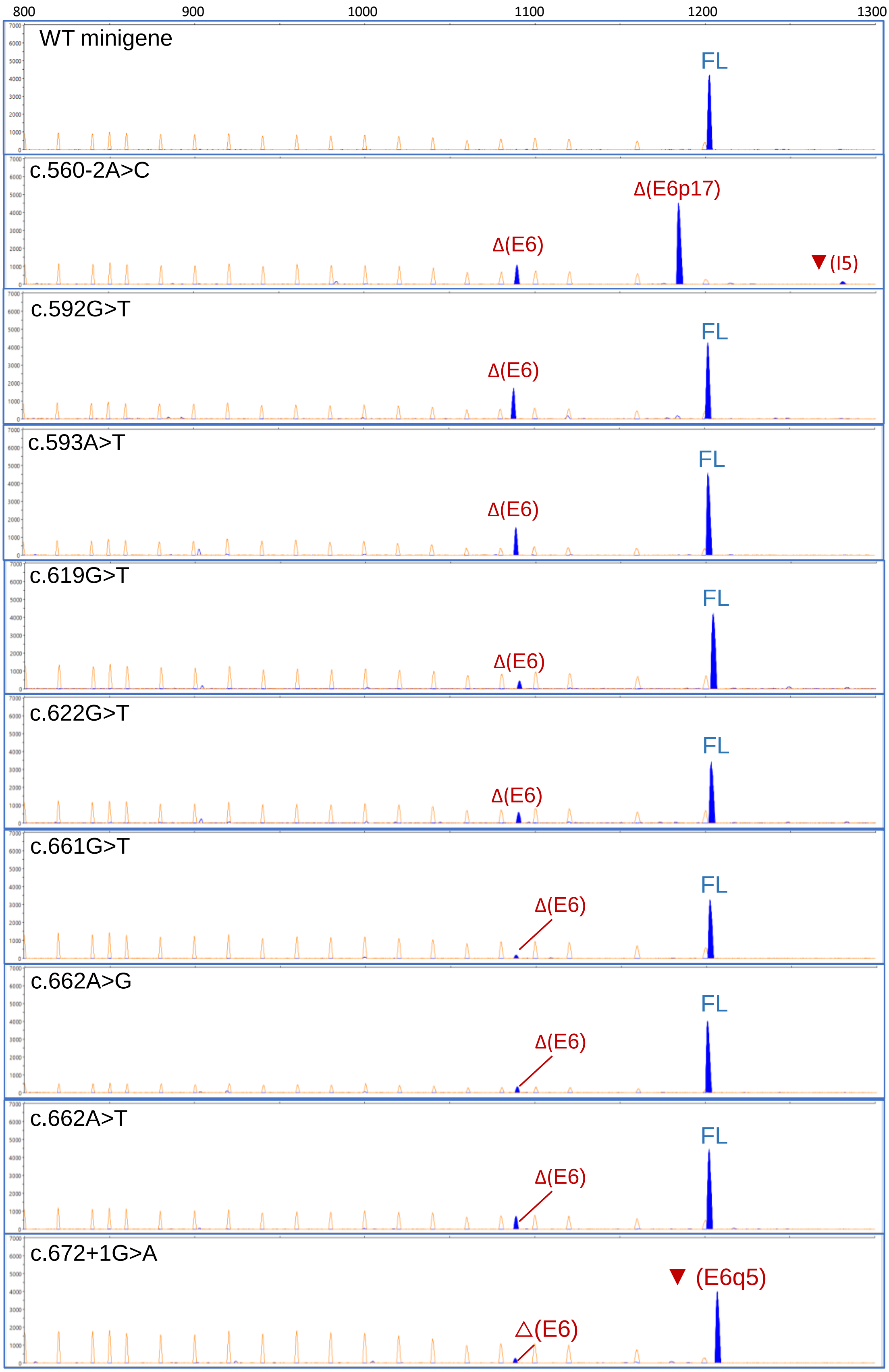
800
900
1000
1100
1200
1300
WT minigene
FL
c.560-2A>C
Δ(E6p17)
Δ(E6)
▼(I5)
c.592G>T
FL
Δ(E6)
c.593A>T
FL
Δ(E6)
c.619G>T
FL
Δ(E6)
c.622G>T
FL
Δ(E6)
c.661G>T
FL
Δ(E6)
c.662A>G
FL
Δ(E6)
c.662A>T
FL
Δ(E6)
c.672+1G>A
▼ (E6q5)
△(E6)

C)

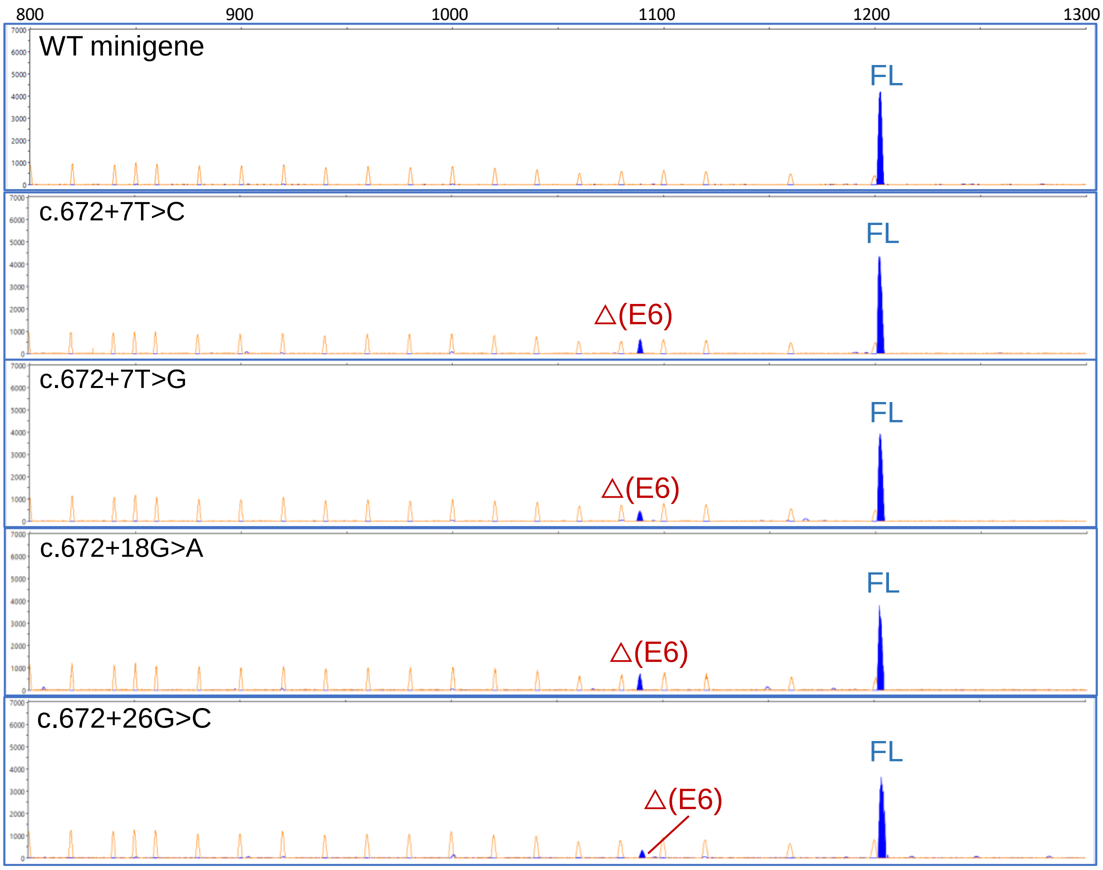

800
900
1000
1100
1200
1300
WT minigene
FL
c.672+7T>C
FL
△(E6)
c.672+7T>G
FL
△(E6)
c.672+18G>A
FL
△(E6)
c.672+26G>C
FL
△(E6)
